# Diatoms vs dinoflagellates: a temporal network analysis of bloom impacts on diversity and phytoplankton community structure in French coastal waters

**DOI:** 10.1101/2025.06.26.661744

**Authors:** Jean-Yves Dias, Victor Pochic, Samuel Chaffron, Pierre Gernez

**Author notes:** <u>Corresponding authors:</u> E-mail address (J-Y. Dias), (S. Chaffron), (P. Gernez).

## Abstract

Understanding phytoplankton community responses to bloom events is essential as these have major implications for biogeochemical cycles, marine food webs, but can also be harmful to ecosystems, the economy, and public health. Diatoms and dinoflagellates are the two most common bloom-forming classes, with distinct ecologies and bloom dynamics. Using time series from a long-term phytoplankton monitoring survey (REPHY) of French coastal waters analysed with ecological association networks and diversity metrics, we investigated how blooms of diatoms and dinoflagellates affect diversity and community structure within phytoplankton communities. We highlight distinct responses: an increase in species richness during diatom blooms and a decrease during dinoflagellate blooms. However, both bloom types resulted in a decrease in alpha diversity indices. Temporal association networks modularity and association strength were also differently impacted between both bloom types. However, high-level association types and composition in association networks remain stable, suggesting an important role of their proportions in ecosystem functioning and resilience. We propose several hypotheses for these findings based on known ecological and biological processes, that remain to be tested individually in dedicated studies. Our work underlines the importance of distinguishing the different types of blooms and studying the role of interactions between phytoplankton for a better understanding of their dynamics.

## Introduction

For decades phytoplankton has been known to play a major role in marine ecosystems as primary producer [1]. It is therefore crucial to develop an advanced understanding of phytoplankton dynamics and particularly bloom events. Bloom studies remain a complex task due to the lack of consensus on what constitutes a bloom. Most studies define blooms as an exceptional biomass, in line with the definition provided by the International Council for the Exploration of the Sea (ICES), which describes blooms as a deviation from the “normal cycle” of phytoplankton biomass [2, 3]. Other definitions rely on thresholds of chlorophyll-a concentration [4] or cell abundance [5], often defined arbitrarily. Some researchers criticize thresholds approaches and argue that the ICES definition is also inadequate for seasonal blooms, as they are part of the “normal cycle” [6].

Several hypotheses have been proposed to understand bloom formation, most of them relying on abiotic conditions [7-10]. More recently, biotic interactions with zooplankton or bacterioplankton have been considered to explain this phenomenon [11-14]. In temperate ecosystems, phytoplankton blooms are expected to occur during spring and to be dominated by large and fast-growing diatoms. It can be followed by shorter summer blooms mainly dominated by smaller diatoms, flagellates, and dinoflagellates [15]. Diatoms and dinoflagellates are the most common bloom-forming classes [15]. Diatoms are generally permanent species with a cosmopolitan distribution. Their blooms are relatively predictable and cyclical. In contrast, dinoflagellates are characterized by short, massive occurrences with high interannual variability. Dinoflagellates display significant ecophysiological diversity, often occupying restricted ecological niches [16]. Some species can dominate the phytoplankton community, forming red tides [17], and be classified as Harmful Algal Blooms due to toxin production or contribution to anoxic conditions [18, 19]. We can therefore expect different impacts of dinoflagellate and diatom blooms on their surrounding communities.

Coastal ecosystems are composed of hundreds of organisms that interact with one another forming complex networks of relationships [20, 21]. More recently, species interactions have gained increasing attention, as they can significantly influence community assembly, species response to climate change, and exacerbate anthropogenic pressures [22, 23]. One debated hypothesis suggests that while the environment selects for functions, biotic interactions select for species [24]. Interactions with viruses, bacteria, and eukaryotic parasites play a crucial role in phytoplankton dynamics [25-28]. Interactions within phytoplankton communities should also be considered. Several studies have highlighted the importance of allelopathy. Allelopathy can have deleterious effects and be considered as a weapon for interspecific competition, or have a positive effect on the whole community if the compounds act against “common enemies” [29, 30]. Competition can also arise among phytoplankton, and some dinoflagellates have developed adaptations such as vertical migrations [31] and mixotrophy [32] to outcompete others. Facilitation is also considered a very common strategy in marine systems [33]. The work of Picoche & Barraquand [34] predicts that phytoplanktonic inter-genus interactions are primarily positive, with competition occurring mainly within genera (strong self-regulation) rather than between different genera. As such, community composition and responses to perturbations are shaped by interactions. While certain interactions may favour the emergence of a bloom, it is likely that blooms themselves exert an influence on the community structure.

Detecting and studying interactions is a complex challenge for ecologists [35]. Association network analyses are commonly used and can be useful to identify associations between species through co-abundance correlations, which can be summarized using graph theory. Distinct community structures will result in different graph typologies that are useful to compare community states and evolution [36, 37]. The present study analyses the impact of blooms on phytoplankton diversity and ecological associations. Using time series from the French phytoplankton monitoring survey (REPHY) analysed through temporal association networks, we addressed the following questions. How do phytoplankton blooms affect the structure and diversity of associations within phytoplankton communities? Are specific types of inter-genus phytoplankton associations favoured during a bloom? Do blooms of diatoms and dinoflagellates have distinct impacts on community structure? Addressing these questions is essential to increase our understanding of bloom impacts on phytoplankton communities and gain deeper insights into differences between diatom and dinoflagellate blooms dynamics.

## Materials and methods

### Phytoplankton and environmental data

The REPHY monitoring survey has been collecting, fortnightly, information on the phytoplankton community through abundances associated with hydrology measurements at the surface since 1987 [38]. Sampling stations are distributed in coastal waters all around mainland France, the sampling and analyses are conducted through a unified procedure [39]. The environmental variables used in this study were seawater temperature, salinity, turbidity, chlorophyll *a* concentration (chl*a*), dioxygen concentration and nutrients concentration: nitrate plus nitrite, ammonia, silicate and phosphate. Phytoplankton samples were analysed following the Utermöhl method [40] using a 10 mL seawater sample. Phytoplanktonic cells with a diameter larger than 20 µm were counted and identified to the lowest taxonomic level. Smaller organisms can also be identified if they were known to be harmful. As a result, REPHY’s definition of phytoplankton is not strictly taxonomic and aligns more closely with microplankton *sensu stricto*. Since different experts identified the phytoplankton, we used the genus as the lowest taxonomic level, to reduce identification bias.

From 195 sampling sites monitored since 1995 we selected 46 sampling stations for which the length of the chl*a* time series without a 2-month gap was longer than 5 years (except during the COVID-19 pandemic). These 46 sampling stations were used to regionalize the dataset (see below). Then, a second selection was conducted to retain 21 chl*a* time series with at least 15 years of data within the same period, i.e., from March 2007 to August 2022 (Fig. 1). Two stations showed an abrupt change in species richness, which we inferred to be due to a change in sampling analysis design. Consequently, these two stations were excluded for further analysis.

**Figure 1:**
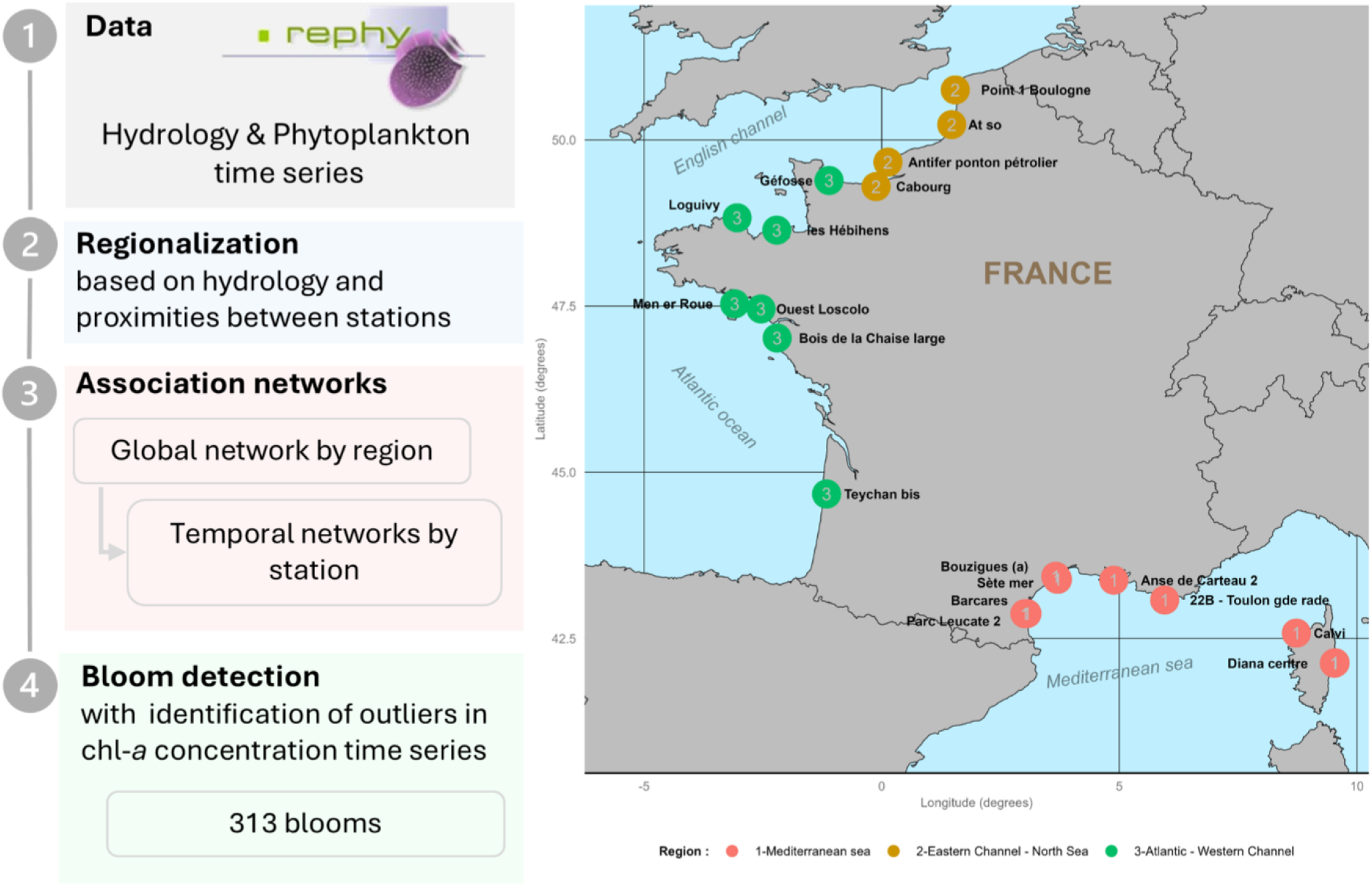
Workflow and map of the sampling sites used in this study. The REPHY sampling sites are coloured by region: Mediterranean Sea (1, red), Eastern Channel – North Sea (2, brown), Atlantic – Western Channel (3, green).

### Hydrology-based regionalization

Our stations cover all mainland French coastlines, which differ in biological and physicochemical ways [41, 42]. To study the broader geographical area beyond each individual station, a hydrology-based regionalization that considers the proximities between them was conducted. This approach enabled us to account for local characteristics that are not captured in the data.

The first step involved handling missing data. To implement this, we used the “EM-PCA” algorithm [43]. In the second step, a principal component analysis (PCA) was performed on all environmental variables. From this PCA, the median position across the dimensions for the 46 stations was calculated. The followings steps were based on the methodology of Chavent *et al*. [44]. Briefly, an initial clustering was carried out using the median PCA positions on all dimensions; a second one was conducted on the spatial locations of the sampling stations using the same method. A geographical constraint weight was defined to maximize hydrological differences while maximizing the spatial constraints. We set this weight to 10% for spatial constraint. A composite distance matrix was generated and used for clustering. The optimal number of regions was determined using the silhouette index.

### Global and temporal association networks

The community structure of phytoplankton was assessed through their biotic associations proxied by their association networks. Association networks are based on correlations between taxa abundances, where nodes represent taxa and edges are associations between pairs of taxa. This approach assumes that correlation implies association. Fourteen graph metrics were computed to capture the main features of the networks (Table 1).

**Table 1:** Definition and abbreviation of the graph metrics used in this study. Italic text provides an ecological interpretation when possible.

| Abbreviation | Definition |
| --- | --- |
| N_nodes | **Number of nodes:** *equal to species richness.* |
| N_edges | **Number of edges:** *number of associations.* |
| Dens | **Density:** corresponds to N_nodes/N_edges. |
| Avg_degree | **Average degree:** average number of edges per node. |
| Diss | **Dissimilarity:** quantifies how well the nodes are connected to each other based on the number of links between them. |
| Trans | **Transitivity:** corresponds to average local connectivity and quantifies how well the adjacent nodes are connected to each other. |
| C_tance | **Connectance:** pourcentage of realized edges out of all possible edges. *Related to association specificity.* |
| Avg_edge_bet | **Average edge betweenness:** quantify the average importance of edges; it is calculated based on the number of nodes that rely on this edge to connect to the hubs (nodes establishing more links with other nodes, which themselves are establishing more). |
| Avg_p_length | **Average path length:** the length of a link is shorter if the correlation between nodes is strong. *Opposite to average associations strength.* |
| Adhes | **Adhesion:** number of edges to delete to disconnect the network. *Related to the sensitivity of the network to the existence of association.* |
| Nat_connect | **Natural connectivity:** number of nodes to delete to disconnect the network. *Related to the sensibility of the network to taxa composition.* |
| Mod | **Modularity:** a set of nodes that have more edges with each other than with the others. *Quantify how the network is organized into sub-communities.* |
| N_clust | **Number of clusters:** number of clusters defined by the fast greedy clustering method. *Rely on identifying sub-communities.* |
| Assort | **Assortativity:** tendency of nodes to connect to nodes with the same number of relationships with other nodes. *If assortativity is positive, this means that highly connected taxa, often generalists that interact with many other species, tend to connect with each other, and similarly for taxa with few connections, specialists.* |

A global association network was performed for each region using phytoplankton abundances. Absent taxa were given an abundance of zero. Only taxa present in more than 30 dates were selected in order to exclude rare taxa and avoid false positive associations. The Spearman correlation coefficient was used to calculate the correlation between taxa abundances. Only significant positive correlations, whose p-value was corrected by the Benjamini-Hochberg method [45], and significantly different to zero, were preserved, using the NetCoMi R package [46].

One of the main limitations of global association networks is their static nature, as they are constructed using the entire time series across all stations within a given region. To deal with temporal dynamics, temporal networks were extracted from the global network. The process involves extracting nodes that correspond to the taxa present at a specific date and station. By doing so, we also kept the associations between pairs of nodes. That means that we have one network per sampling date for each station, allowing us to calculate graph metrics for each temporal network as well as for the global network. To reduce the dimensionality of the different graph metrics, a PCA was performed (Fig.3).

### Network compositions

To analyze association networks, we first examined their taxonomic composition, determined by the number of nodes belonging to each taxonomic class. We focused on 3 classes: diatoms (*Bacillariophyceae*), dinoflagellates (*Dinophyceae*) and others (“Other phyla”). We also analyzed the composition of edges, which provides insights into the “association types” within the network. We focused on: association between diatoms, between dinoflagellates, between diatoms and dinoflagellates, and any association involving a taxon from other phyla.

The analysis of the association network compositions provides the proportion of each association type. To determine whether these proportions result from ecological processes rather than basic taxonomic composition, randomization tests were performed. If the proportions were solely due to composition, we would expect random associations to yield similar proportions. However, while it is well-established that associations themselves are influenced by ecological processes, it is less clear whether this also applies to the association type proportions. To assess this, we took the association matrix for each region. Simple random permutations were performed; by doing so, species richness and the number of associations remained the same. This process allowed us to calculate the proportion of each association type in a “null” association network. 10 000 null association networks were generated for each region. We compared the null distribution of the proportion of an association type to the observed proportion of that association type in the global network by calculating p-values. Because we obtained a p-value for each association type, a global p-value was calculated based on Fisher’s correction for each region.

### Diversity indices

Three diversity indices were calculated from the abundance data for each sampling day. The Shannon index is a biodiversity index that considers species richness and equitability [47]. The Piélou index is used to estimate how equitable the abundances of species in a community are [48]. The Berger-Parker index corresponds to the proportion of the most abundant taxon to estimate its dominance [49].

### Bloom detection and characterization

The definition of bloom for our study is based on the ICES definition considering the comments of Isles & Pomati [6]: a bloom is a deviation to the “normal” seasonal cycle of phytoplankton biomass. Here, we used chl*a* as proxy of biomass. The time series for each station were processed equally. Chl*a* time series were regularized using linear interpolation. The seasonal component was subtracted by seasonal-trend decomposition by Loess [50] assuming that the seasonal component is identical over years. Phytoplankton blooms were identified from the deseasonalized chl*a* time series using outlier detection through the median absolute deviation [51]. The thresholds coefficients described in Leys et al. [51] were chosen to be the more objective.

For each bloom detected we assumed that the most abundant taxa were responsible for it. We determined the first previous sampling date as the “before” moment and the first following sampling date as the “after” moment of the bloom. If there are successive dates that are determined as blooms and if the class remains the same (*e*.*g*., from *Chaetoceros* to *Skeletonema* blooms, two diatoms), we consider only one “before” and “after” moments. If there are successive dates that are determined as blooms but the class changes (*e*.*g*., from diatom to dinoflagellate bloom), we consider the first bloom with a “before” moment and the second with an “after” moment. If a sampling date is both an “after” and a “before” we excluded it from bloom analysis.

To estimate whether a bloom means a higher dominance of a particular taxon, we compared the Berger-Parker index for bloom and “non-bloom” dates. For this analysis, 100 random “non-bloom” dates were selected, controlling for the season (spring and summer are overrepresented within bloom dates). We compared this across all regions and by region, using a Wilcoxon test.

### Statistical analysis and software

Statistical significance was defined as p < 0.05 after correction for multiple testing. Kruskal-Wallis tests were used to assess differences between regions, types of blooms, and bloom moments. If the test was significant, a Dunn test with Benjamini-Hochberg’s method for p-value correction was done. All analyses were performed using R 4.4.2 [52]. Further details are available in the Data availability section.

## Results

### Phytoplankton associations and diversity in French coastal waters

Three regions were obtained from the regionalization process (Fig. 1). The first region corresponds to the Mediterranean Sea, the second to the Eastern Channel – North Sea (hereafter Eastern Channel-NS), and the third to the Atlantic Ocean – Western Channel (Atlantic-WC).

The mean relative abundance of diatoms was 72.10% in the Mediterranean Sea and 75% for the other two regions. Dinoflagellates are the second most dominant class, with a mean proportion of 17.69% in the Mediterranean Sea and approximately 6% in the other regions. The three regions differed in terms of phytoplankton composition at the genus level (Fig. 2). For example, there was a higher relative abundance of *Skeletonema* in the Mediterranean Sea, *Phaeocystis* in the Eastern Channel-NS and *Asterionnellopsis* in the Atlantic-WC region. Analysing the temporal pattern of composition revealed that extreme dominance of the community by one taxon can display a well-established seasonality (for instance, *Phaeocystis* dominance occurs consistently around March in the Eastern Channel-NS) or emerge suddenly (for instance, a *Nitzschia* dominance was observed in July 2019 in the Atlantic-WC).

**Figure 2:**
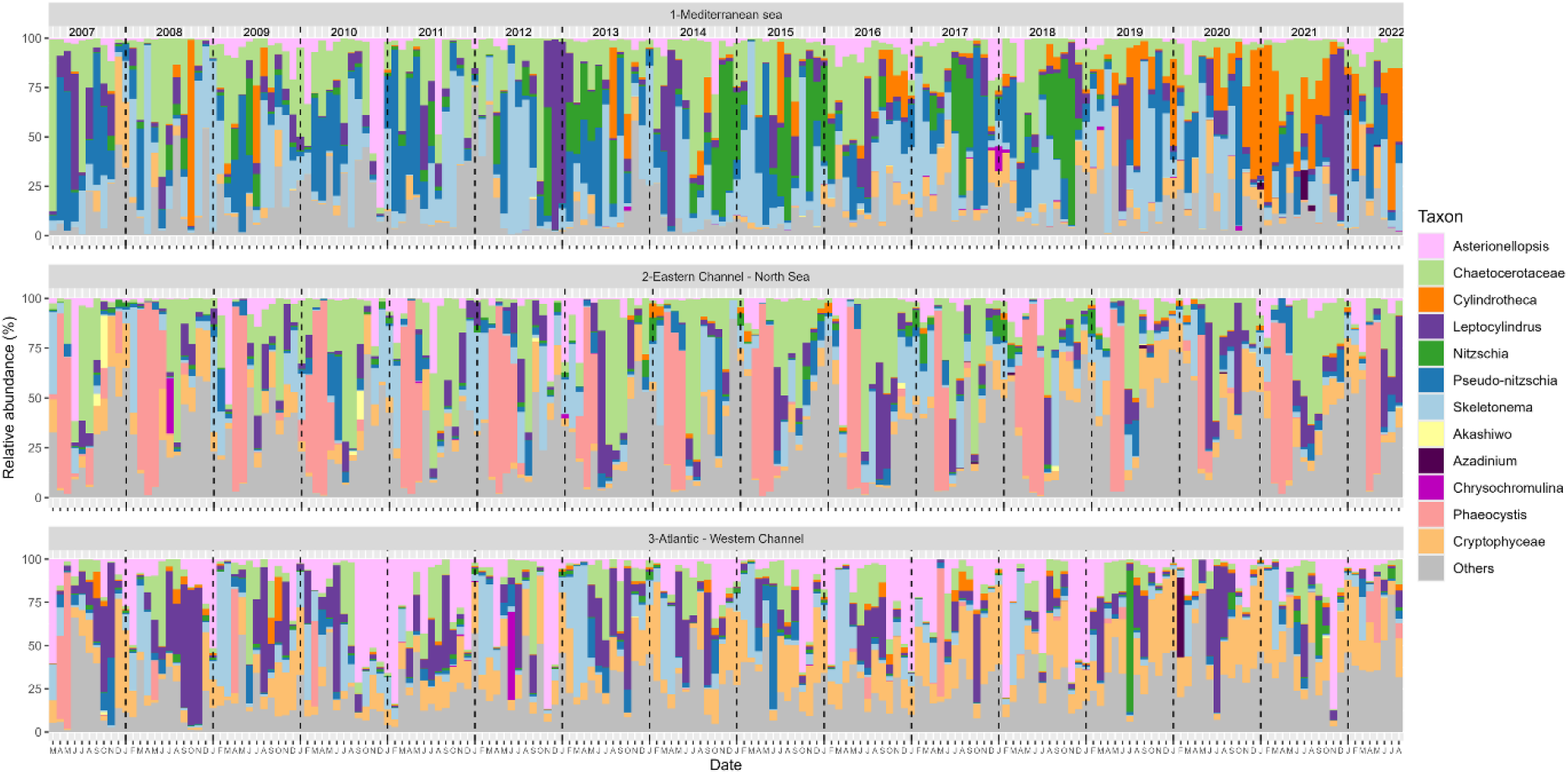
Average relative abundance by month and region from 2007 to 2022 for the four most abundant taxa by season and region.

The PCA based on the graph metrics allows us to identify the association network structures and also how graph metrics are related to each other (Fig. 3). The first three dimensions of the PCA explained 74% of the total variance in graph metrics (38.5%, 24.7% and 10.8%, respectively). The first dimension (Dim1) primarily captures the variations in species richness and the robustness of the network. Along this dimension, the number of nodes, edges, density, and adhesion are positively correlated with each other. The second dimension (Dim2) can be interpreted in terms of modularity and association selectivity. Along Dim2, modularity and dissimilarity are positively correlated with each other, while connectance and transitivity negatively correlated with Dim2. The third dimension is mainly positively correlated with average path length, meaning that high Dim3 values indicate a low average association strength.

**Figure 3:**
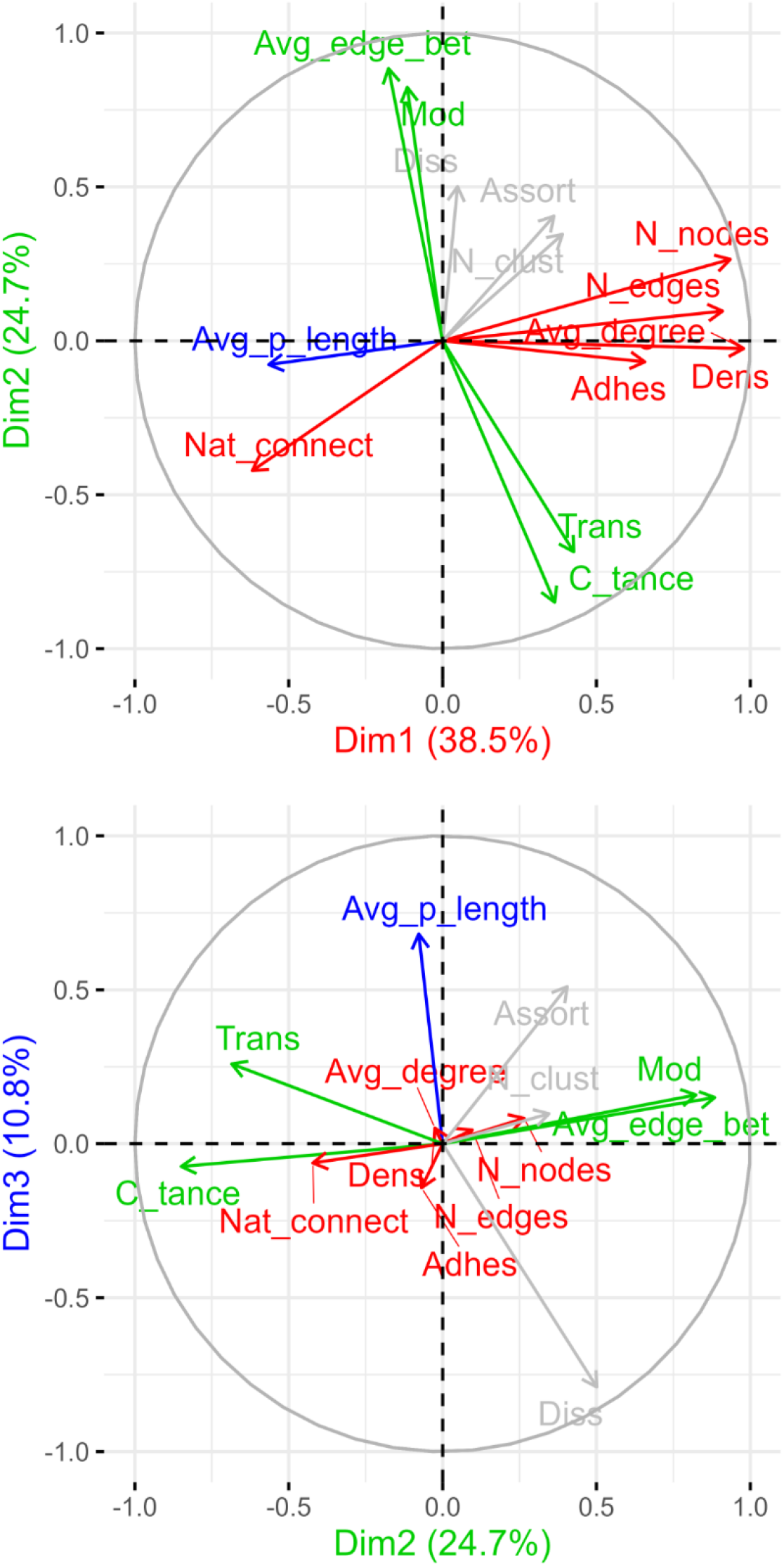
Principal Component Analysis (PCA) based on graph metrics. The variance explained by each dimension is indicated in the axis labels. Graph metrics were calculated for all temporal networks across all regions. See Table 1 for the definition of each abbreviation.

Regional global association networks differed by structure and association types (Fig. 4). The Mediterranean Sea region has the lowest species richness and is more sensitive to the extirpation of a single species (Nat_connect) (Fig. S1). Conversely, the Atlantic-WC region has the most robust network and was significantly more organized into sub-communities with high association specificity than the Eastern Channel-NS, which was, in turn, more structured than the Mediterranean Sea region. The average association strength was significantly lower in the Eastern Channel-NS region, while it was similar between the other two regions (Fig. S1). Most associations were positive: 75.97%, 63.76% and 64.46% in the Mediterranean, Eastern Channel-NS and Atlantic-WC networks, respectively. In accordance with phytoplankton counts, diatoms represented the largest proportion of taxa in the global networks, followed by dinoflagellates (Fig. 4). Consistently, associations between diatoms were the most frequent across all regions, followed by associations between diatoms and dinoflagellates, except in the Atlantic-WC region where 31.6% of the links were related to other types of associations (Fig. 4). It is important to remember that this applies to the global networks, and there is a great temporal variability (Fig. S2).

**Figure 4:**
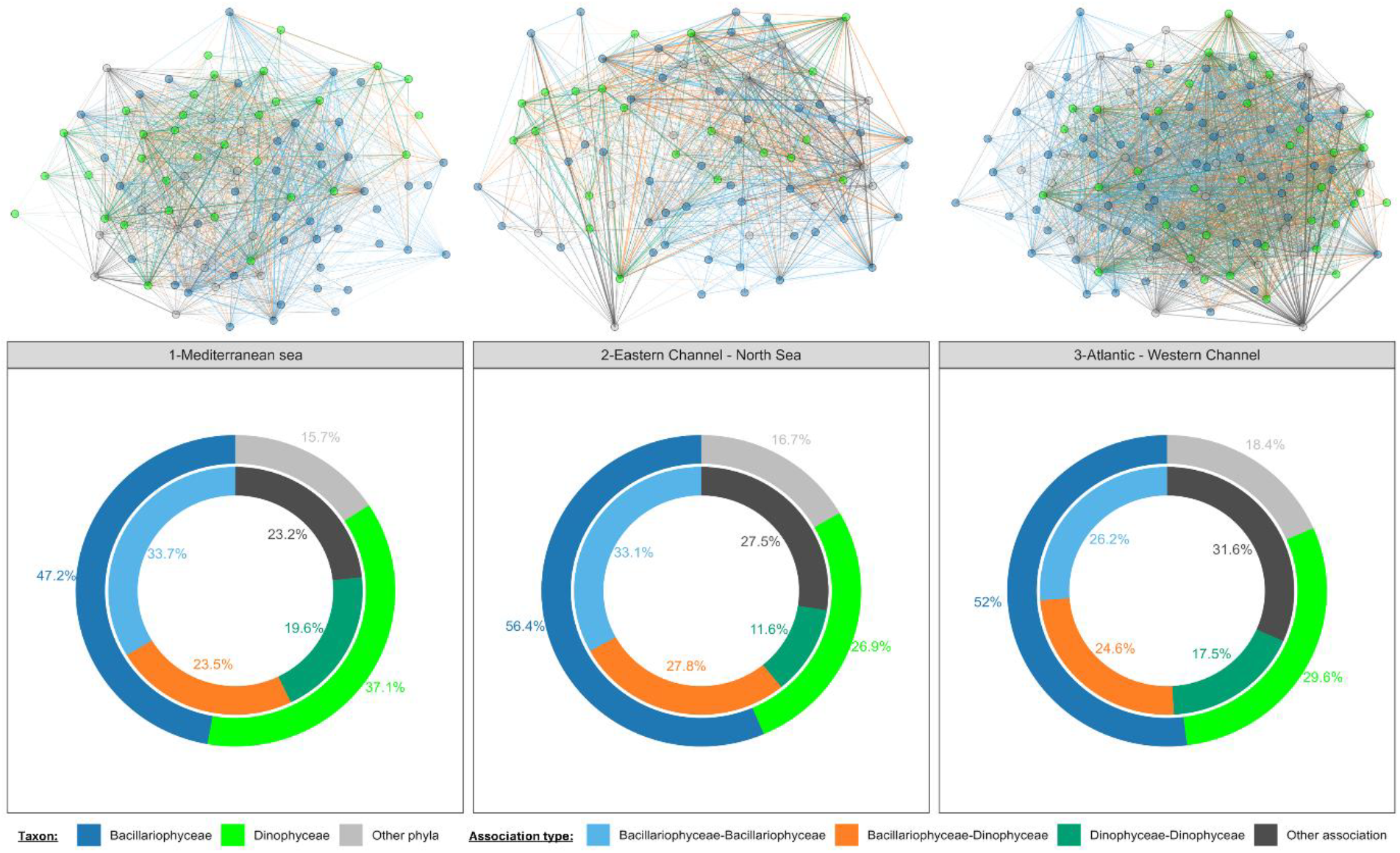
Global association networks and their composition by region. In the global association networks (top), nodes represent *Bacillariophyceae* (dark blue), *Dinophyceae* (green), or other phyla (grey). Links are coloured according to the type of association: associations between *Bacillariophyceae* (light blue), between *Dinophyceae* (dark green), between *Bacillariophyceae* and *Dinophyceae* (orange), or other types of associations (dark). Thicker links indicate stronger associations. In the donut plots (lower part), the outer circle represents the relative number of each taxon in the global network, while the inner circle indicates the relative proportion of each association type. The colour legend is consistent with that of the global association networks.

Importantly, the proportion of association types results from ecological processes and not from random processes. In all regions, the association type proportions significantly differed from a random distribution of associations: (the randomization test had a p-value = 0 in the Mediterranean Sea (i.e., some observed proportions were never found in the random distribution), a p-value = 2.34.10^−10^ in the Eastern Channel-NS, and a p-value = 0 in the Atlantic-WC (Fig. S3).

### Bloom detection

Our method for bloom detection identified 125 blooms in the Mediterranean Sea, 67 in the Eastern Channel-NS, and 121 in the Atlantic-WC region. Two-thirds of all detected blooms were blooms of diatoms (Table 2; Fig. S4). Cryptophytes in the Mediterranean Sea, haptophytes in the Eastern Channel-NS, and dinoflagellates in the Atlantic-WC region were the second most frequent classes in terms of bloom occurrence. 13 dinoflagellate blooms were detected in the Mediterranean Sea, 5 in the Eastern Channel-NS, and 10 in the Atlantic-WC region. Additionally, 14 haptophyte blooms related to *Phaeocystis* were identified in the Eastern Channel-NS. These small number of blooms may limit the statistical power to evaluate their impacts. The genera responsible for blooms differed between regions (Table 2). For example, 10 *Cylindrotheca* blooms were detected in the Mediterranean Sea, of which 9 in coastal lagoons, but none in the other regions. *Prorocentrum* blooms were the most detected dinoflagellate blooms in the Mediterranean Sea whereas it was *Lepidodinium* in the two other regions.

**Table 2:**
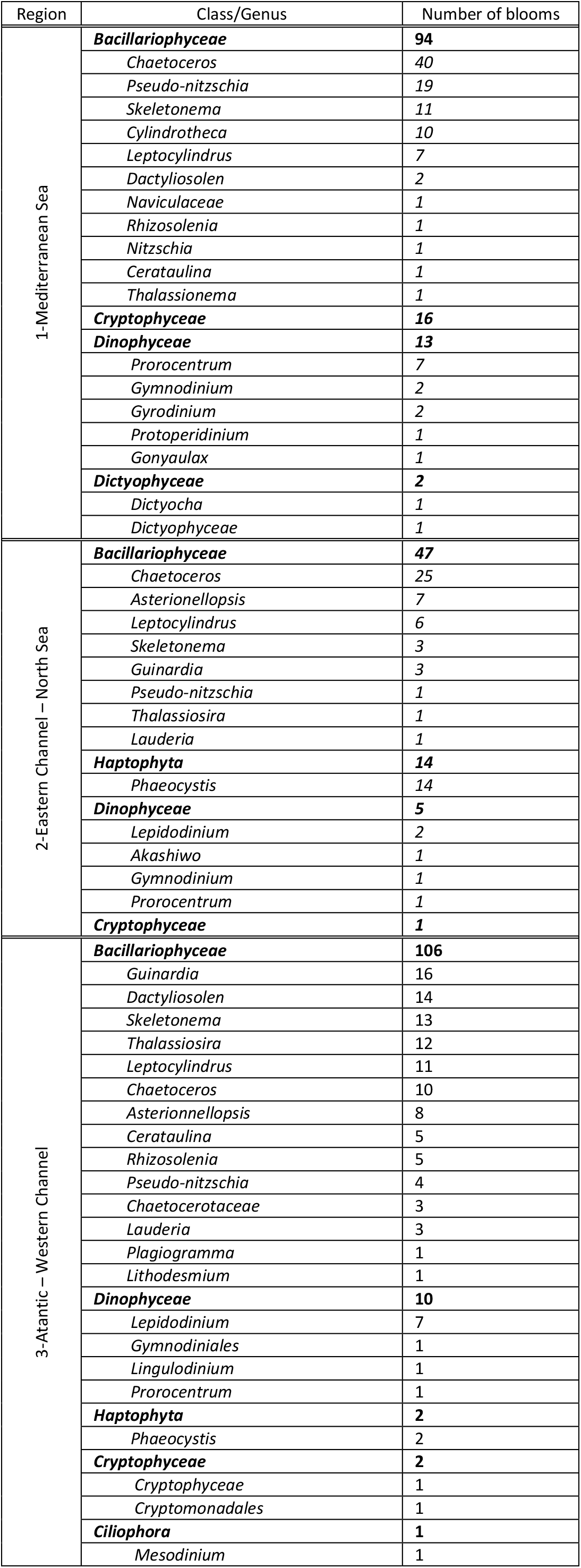
Blooms detected by region, categorized by class and genus.

| Region | Class/Genus | Number of blooms |
| --- | --- | --- |
| 1-Mediterranean Sea | **Bacillariophyceae** | **94** |
|  | *Chaetoceros* | 40 |
|  | *Pseudo-nitzschia* | 19 |
|  | *Skeletonema* | 11 |
|  | *Cylindrotheca* | 10 |
|  | *Leptocylindrus* | 7 |
|  | *Dactyliosolen* | 2 |
|  | *Naviculaceae* | 1 |
|  | *Rhizosolenia* | 1 |
|  | *Nitzschia* | 1 |
|  | *Cerataulina* | 1 |
|  | *Thalassionema* | 1 |
|  | **Cryptophyceae** | **16** |
|  | **Dinophyceae** | **13** |
|  | *Prorocentrum* | 7 |
|  | *Gymnodinium* | 2 |
|  | *Gyrodinium* | 2 |
|  | *Protoperidinium* | 1 |
|  | *Gonyaulax* | 1 |
|  | **Dictyophyceae** | **2** |
|  | *Dictyocha* | 1 |
|  | *Dictyophyceae* | 1 |
| 2-Eastern Channel – North Sea | **Bacillariophyceae** | **47** |
|  | *Chaetoceros* | 25 |
|  | *Asterionellopsis* | 7 |
|  | *Leptocylindrus* | 6 |
|  | *Skeletonema* | 3 |
|  | *Guinardia* | 3 |
|  | *Pseudo-nitzschia* | 1 |
|  | *Thalassiosira* | 1 |
|  | *Lauderia* | 1 |
|  | **Haptophyta** | **14** |
|  | *Phaeocystis* | 14 |
|  | **Dinophyceae** | **5** |
|  | *Lepidodinium* | 2 |
|  | *Akashiwo* | 1 |
|  | *Gymnodinium* | 1 |
|  | *Prorocentrum* | 1 |
|  | **Cryptophyceae** | **1** |
| 3-Atlantic – Western Channel | **Bacillariophyceae** | **106** |
|  | *Guinardia* | 16 |
|  | *Dactyliosolen* | 14 |
|  | *Skeletonema* | 13 |
|  | *Thalassiosira* | 12 |
|  | *Leptocylindrus* | 11 |
|  | *Chaetoceros* | 10 |
|  | *Asterionellopsis* | 8 |
|  | *Cerataulina* | 5 |
|  | *Rhizosolenia* | 5 |
|  | *Pseudo-nitzschia* | 4 |
|  | *Chaetocerotaceae* | 3 |
|  | *Lauderia* | 3 |
|  | *Plagiogramma* | 1 |
|  | *Lithodesmium* | 1 |
|  | **Dinophyceae** | **10** |
|  | *Lepidodinium* | 7 |
|  | *Gymnodiniales* | 1 |
|  | *Lingulodinium* | 1 |
|  | *Prorocentrum* | 1 |
|  | **Haptophyta** | **2** |
|  | *Phaeocystis* | 2 |
|  | **Cryptophyceae** | **2** |
|  | *Cryptophyceae* | 1 |
|  | *Cryptomonadales* | 1 |
|  | **Ciliophora** | **1** |
|  | *Mesodinium* | 1 |

### Blooms impact on diversity, community structure and associations

Blooms impact phytoplankton diversity: on average, the Shannon and Piélou indices were lower during dinoflagellate or diatom blooms, except in the Eastern Channel-NS region, which exhibited a different pattern (Fig. 5A). The Berger-Parker index generally increased during blooms. Coherently, when comparing bloom to non-bloom samples, our results indicate a higher Berger-Parker index (12%), which seems to be mainly driven by the Mediterranean region (Fig. S5). Additionally, no correlation between chl*a* concentration and the Berger-Parker index (Spearman’s correlation) was found.

**Figure 5:**
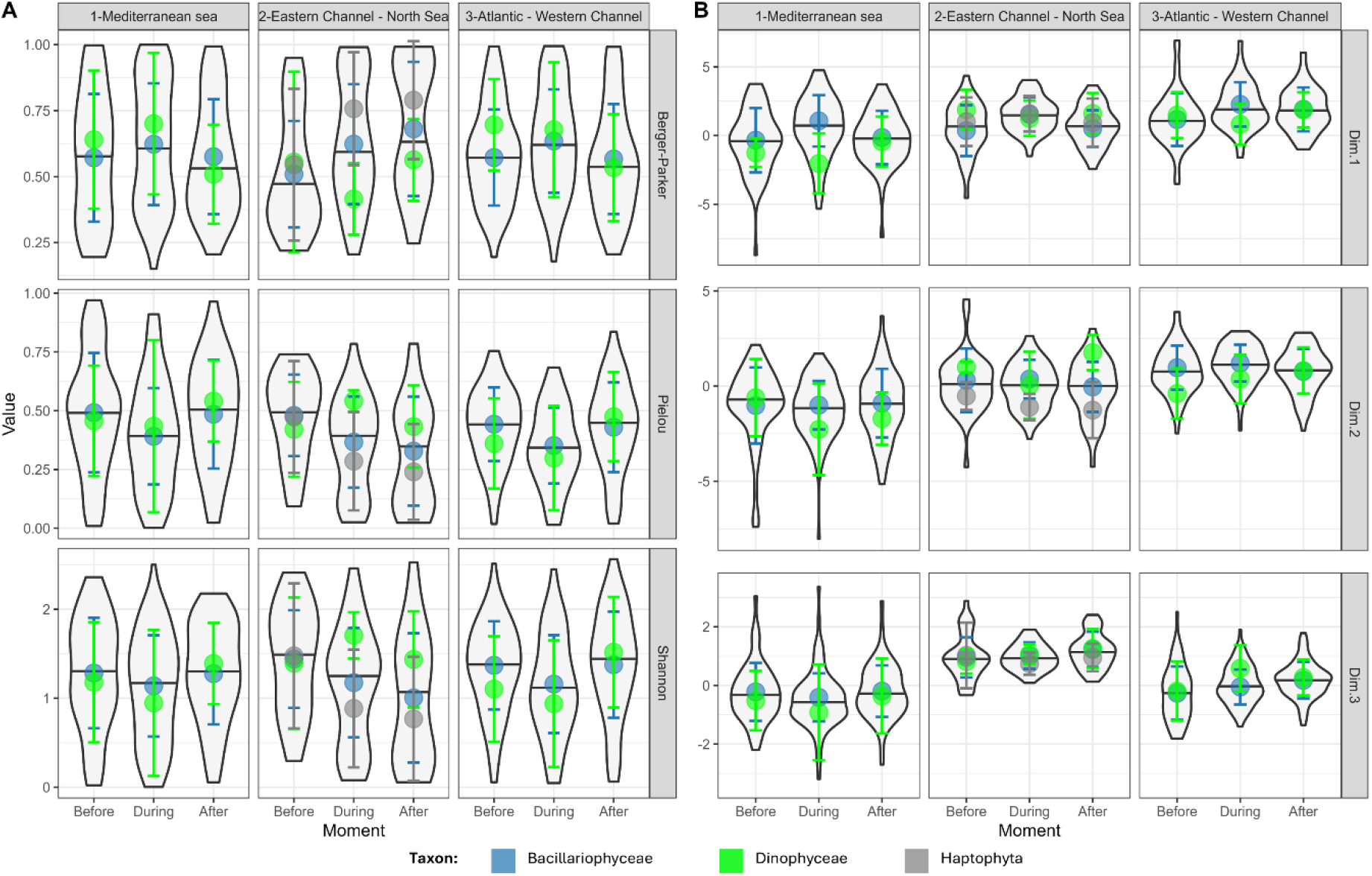
Diversity indices (A) and PCA dimensions (B) before, during, and after a bloom for each region. The violin plot shows the distribution of values for diatom and dinoflagellate blooms, as well as Eastern Channel’s haptophyte blooms. The black line represents the mean value. Points indicate the mean value for *Bacillariophyceae* (dark blue), *Dinophyceae* (green), and *Haptophyta* (grey) blooms. Error bars represent the standard deviation.

Blooms also impact the structure of association networks (Fig. 5B). Our results revealed an increase in the Dim1 PCA coordinates during diatom and haptophyte blooms, with varying amplitudes depending on the region. In contrast, during dinoflagellate blooms, Dim1 values decreased. These findings suggest that diatom and haptophyte blooms are associated with an increase in species richness and network robustness (Adhesion), whereas dinoflagellate blooms have the opposite effect. Blooms’ impacts on modularity and association strength appeared minimal.

Blooms maintain the proportions of taxonomic composition in the networks (Fig. 6). Our results indicate that during blooms, the relative proportions of taxa in the association networks remained stable. While species richness changed (Fig. 5B), no taxonomic class exhibited a significant increase in proportion at any moment of the bloom (Fig. 6). This indicates a stability in the taxonomic composition of association networks. Although differences emerged depending on the bloom taxonomic class (e.g., proportion of dinoflagellates was higher whatever the phase of dinoflagellate blooms compared to other bloom types), this is probably due to different seasonal preferences between classes.

**Figure 6:**
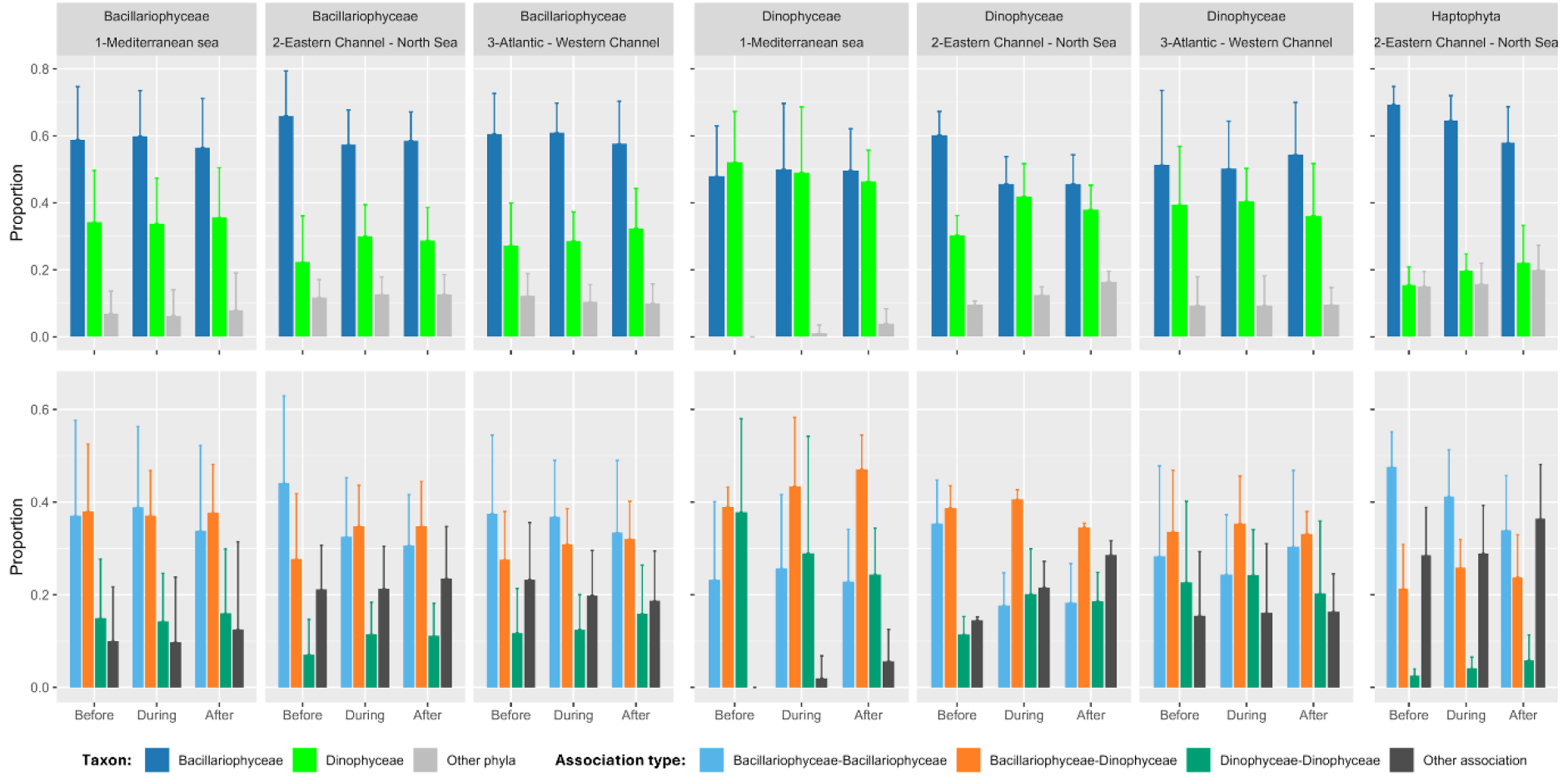
Composition of association networks before, during, and after *Bacillariophyceae, Dinophyceae*, or *Haptophyta* blooms for each region. The top bar plots show the average proportion of each taxon on the association networks (*Bacillariophyceae*, dark blue; *Dinophyceae*, green; other phyla, grey). The bottom bar plots display the average proportion of each association type (between *Bacillariophyceae*, light blue; between *Bacillariophyceae* and *Dinophyceae*, orange; between *Dinophyceae*, dark green; and other types of associations, dark). Error bars represent the standard deviation.

A regional similarity was observed for each bloom type, though it was less consistent for dinoflagellate blooms, especially for the ‘Other association’ type in the Mediterranean. Examining the networks association types revealed that, like taxonomic composition, proportions remained relatively stable across bloom phases, and regions, while the proportion of association types varied depending on the bloom taxonomic class (Fig. 6). Around dinoflagellate blooms, networks contained more associations involving dinoflagellates than other bloom types, while around haptophyte blooms, “Other association” types were more prevalent.

Overall, these findings highlight that bloom impacts depend on the taxonomic nature of the dominant taxon, but also reveal similarities across regions regarding a general stability of association networks in taxonomic and association-type composition across different bloom phases.

## Discussion

### Association networks to study phytoplankton communities

Association network analysis is useful to estimate associations between taxa. Here, “association” does not necessarily mean “interaction” as we cannot disentangle both nor define the type of interaction. Different interactions can result in the same signal (*i*.*e*., facilitative or mutualistic). It is likely that correlations do not reflect only biotic interactions but also largely overlapping niches. Another limitation of this approach is that it provides a global association coefficient. This implies that we cannot consider changes in the signs or strength of the association between two given taxa. Indeed, species may modify their interactions in the presence of others or in different environmental conditions [35]. This modification can have a great impact on the overall ecosystem structure and functioning [53]. Previous modelling studies demonstrated that interaction strength variability is a key component for ecosystem stability and species coexistence because species interactions can be modulated by biotic or abiotic changes [54, 55]. Despite such limitations, the relevance of using association networks to study phytoplankton interactions has been demonstrated: such approach was proven to be able to recover known interactions as well as discover new ones [56, 28].

Our results provide an understanding of what association networks can capture in terms of community ecology. The number of nodes, edges, and adhesion covary. This can be explained because the number of edges follows a quadratic function with the number of nodes [57]. Modularity increases when connectance decreases, meaning that the community is more organized into sub-communities when association specificity is high. It is interesting to note that species richness is not related to sub-community organization, indicating that an additional taxon can be well integrated into a sub-community. Studies have found a positive correlation between robustness and connectance, which is not supported by our analysis [58, 59]. Another important graph metric is the average association strength (Dim3). We initially hypothesized that weak association strength could indicate a more sensitive community because taxa are less influenced by each other. However, the work of Allesina & Tang [60] supports the idea that stability is achieved by decreasing the average interaction strength, thereby lowering complexity. Accordingly, Carpentier et al. [57] demonstrated a trade-off between robustness and stability in biological networks. Other work from Thebault and Fontaine [61] concluded that stability of trophic networks is higher in modular and weakly connected architectures. Altogether, it appears that community stability is difficult to estimate through graph metrics in our analysis, since robustness, modularity and interaction strength are not linked.

### Phytoplankton communities and association structures in French coastal waters

Our regionalization process identified three coastal regions in France. It is well known that the Mediterranean Sea is a distinct ecosystem compared to the Atlantic Ocean and the English Channel [62, 42], accentuated in our dataset by the prevalence of coastal lagoons in the Mediterranean region. This regionalization may seem trivial, but we argue that it is necessary to differentiate regions objectively and without prior assumptions. The fact that the Western Channel stations are more similar to the Atlantic stations supports this approach.

The three regions presented distinct communities, consistent with their different local environments and ecosystem functioning. The Shannon index was lower in the Mediterranean Sea, which could be partly explained by species smaller than 20 µm that are not systematically counted [63]. Diatoms were found to be dominant in phytoplankton communities across all regions, which is consistent with previous findings [64]. In all regions, most associations were positive, a result also observed in other studies [13, 28, 65]. This supports the idea that inter-genus associations are primarily positive as argued by Picoche & Barraquand [34]. Different association structures were found between regions, reflecting distinct communities. The Atlantic-WC region appears to have the highest species richness and to be more organized into sub-communities. However, this may be influenced by the higher number of sampling dates in this region, potentially introducing biases and apparently increasing the number of detected associations (*ie*., sampling effect; [66, 67]).

A key finding of our study is that association type proportions result from ecological processes, and therefore they must be further investigated. This implies association type proportions likely play a key role in community equilibrium and responses to perturbations. The diversity of interaction types between species remains understudied, as they are difficult to capture, predict and validate. Previous works [68, 69] demonstrated that the balance of interaction types plays an important role in community dynamics. Thus, the diversity and proportion of interaction types may also play a crucial role in community responses to perturbations and overall ecosystem functioning.

### Blooms impact on diversity

The method used for bloom detection has several advantages: it is simple, auto-adaptive to the study site, and does not rely on prior assumptions. Defining the phases of a bloom as the first previous and following dates allows for analysis of bloom dynamics without considering temporal changes in abiotic factors. Blooms occurred in all seasons: this supports the idea that studying blooms all year long (and not only during spring) is essential for a better understanding of phytoplankton ecology, as previously highlighted by Smayda [3]. We detected a relatively large number of blooms, with most blooms being dominated by diatoms, as generally observed in coastal waters [15, 64]. Dinoflagellate blooms may be less frequent, shorter and/or more localized, and therefore less likely to be sampled than blooms of diatoms [18].

A key result is that species richness increases during diatom and haptophyte blooms but decreases during dinoflagellate ones. It suggests different community responses during these blooms. Several hypotheses can explain lower species richness during dinoflagellate blooms. Some dinoflagellates have deleterious effects on other phytoplankton, excluding competitors through allelopathy [29, 70] or predation for mixotrophy [71, 72]. Beyond direct interactions, environmental conditions favouring their blooms may be less suitable for diverse phytoplankton taxa. For instance, in heavily stratified waters, intense vertical migrations allow dinoflagellates to exploit deep, nutrient-rich layers inaccessible to their competitors [31]. This aligns with the observation that dinoflagellate blooms occur more sporadically and ephemerally than diatom blooms, with more restrictive ecological niches [16]. Contrary to dinoflagellate blooms, diatom and *Phaeocystis* blooms showed higher species richness than before and after the bloom. During *Phaeocystis* blooms, cells can agglomerate and form mucus, which acts as a defence against grazers [73, 74]. Reduced grazing pressure may enhance species richness. Diatom blooms may exhibit greater species richness due to the facilitative actions of taxa and better self-regulation, allowing them to coexist. However, all these hypotheses remain to be tested using mesocosms experiments, as we cannot directly assess the underlying processes. Investigating the ratio of positive to negative associations during dinoflagellate and diatom blooms, as well as their dynamics, could provide further insights. The abiotic conditions that favour each type of bloom remain a key challenge in marine ecology [75].

Blooms also impact diversity. The Berger-Parker index increased during blooms, regardless of chl*a* levels, highlighting the limits of some bloom definitions based on abundance thresholds. This pattern suggests dominance by a single taxon, likely due to competitive exclusion mechanisms such as allelopathy, mixotrophy, nutrient affinity or grazing resistance [29, 32, 70, 76-78]. In 2 regions (Mediterranean and Atlantic), the Shannon and Piélou indices decreased during blooms, reflecting uneven species abundances and a community dominance, as confirmed by the Berger-Parker index. However, in the Eastern Channel-NS region, the diversity indices have a different pattern, likely due to specific bloom-forming taxa, abiotic environment, or community structure. These diversity changes were not linked to species richness, indicating that blooms reorganize the abundances rather than change species presence. This highlights the limitation of alpha diversity metrics and supports the use of abundance distribution analysis for a more complete overview [79, 80].

### Blooms impact on community structure and associations

The impact of blooms on community structure is more difficult to disentangle as our results do not give a clear signal regarding graph metrics. Additional higher resolution data could reveal effect of blooms on community structure and associations. In that case, the high variability observed would indicate that these effects cannot be adequately studied with a classification as broad as all diatom or dinoflagellate blooms, regardless of season, and separated into three regions. This issue brings us back to a fundamental challenge in ecology: the need to group elements into broader categories to understand certain processes, even when some phenomena may not be fully captured within these broader groupings [81, 82].

### The importance of equilibria in phytoplankton blooms

A key result is that broad taxonomic composition and types of association remain unchanged during blooms, regardless of bloom type or region, suggesting that no association type is favoured during a bloom. A surprising finding is that the composition and proportions of associations are highly similar across regions. Our randomization test confirmed that association type proportions are shaped by ecological processes and not merely a reflection of the proportion of each taxonomic class (Fig. S3). The fact that species richness changes during blooms makes this result even more unexpected. Blooms may not represent a complete renewal of the community but rather a reorganization of abundances through self-regulation and compensatory mechanisms. Jochimsen et al. [83] and Gonzalez & Loreau [84] demonstrated that compensatory dynamics and interaction types are key factors contributing to phytoplankton resilience and stability. This aligns with the idea that association type proportions are essential for the resilience and functioning of phytoplankton communities through regulatory mechanisms. This also supports the idea that competition primarily occurs within genera [34]. The community persisting during blooms may consist of taxa with high functional redundancy relative to the “non-bloom” community. In addition, functional redundancy is known to enhance community resilience [85-87]. Taxonomic classification overlaps significantly with functional classification, further supporting this argument [88, 89]. Exploring the notion of a core community could be particularly insightful. A core community consists of taxa that consistently occur, are locally abundant, and regionally common, in contrast to satellite or transient communities, which are sparse and occur less frequently [90-92]. Core species are hypothesized to be better adapted to local environmental conditions than transient species, and therefore, more likely to exhibit traits indicating environmental tolerance [91]. The core community may remain relatively stable, as observed in our analysis, while transient species are more affected by blooms. Consequently, the core community could exhibit distinct characteristics in terms of modularity, species richness, and average association strength. Xue et al. [93] demonstrated, through network analysis, that core communities exhibit more stable structures against perturbations than satellite communities. Blooms are likely not disruptive enough to impact this core community, which ensures resilience and a return to the previous functional state. However, methodological challenges and ongoing debates persist regarding the validity of this classification [94, 95]. Overall, the proportions differ between diatom, dinoflagellate, and haptophyte blooms. This highlights that when diatoms bloom, there are proportionally more diatoms both before and after the bloom, along with a higher number of associations involving diatoms, likewise during dinoflagellate blooms: more dinoflagellates and more associations involving them. This result suggests that the environmental context and abiotic forcing play a key role in shaping both community composition and the proportions of association types, more so than the influence of blooms themselves. Integrating bacteria and viruses could provide insightful information to unravel the full complexity of blooms [96].

## Conclusion

Through the analysis of 16 years of phytoplankton abundance data using association networks, our study highlighted the influence of blooms on phytoplankton community structure, species richness, and association dynamics. We show that diatoms and dinoflagellates reduce diversity indices, likely due to a reorganization of abundances rather than competitive exclusion. On the other hand, diatom and haptophyte blooms increase species richness, whereas dinoflagellate blooms lead to its decline. Despite these variations, taxonomic composition and association type proportions remain unchanged during blooms across all regions. This suggests strong regulatory mechanisms within phytoplankton communities and a role of these proportions in community functioning. In addition, there seems to be no specific types of inter-genus phytoplankton associations favoured during a bloom. Diatoms and haptophytes respond to bloom conditions in a more similar way, whereas dinoflagellates generally show opposite patterns. This supports the idea of distinct mechanisms initiating and supporting these types of blooms. Future studies should refine these findings by integrating bacteria and virus samplings, functional classifications, exploring the role of sub- or core-communities, and testing precise ecological processes. Our work confirms the importance of studying biological interactions to understand bloom dynamics at a community level.

## Data availability

The raw data that support the findings of this study were provided by IFREMER and available from SEANOE, at https://doi.org/10.17882/47248. All R scripts and generated data used in this study are available from the Github repository, at github.com/J-YDi/Diatoms-vs-Dinoflagellates and from zenodo at 10.5281/zenodo.15516291.

## Acknowledgments

This study was carried out thanks to the long-term work of all past and present members of the IFREMER REPHY monitoring network; from managers, vessel crews to phytoplankton analysers people, authors would like to warmly thank each of them. Authors thank Eric Goberville (BOREA, Sorbonne University) and François Lantoine (LECOB, Sorbonne University) for their relevant comments on a previous version of this work. Authors thank the French government and taxpayers that, still, support oceanographic research financing the REPHY network and our salaries.

## Author contributions

JYD, VP, PG, and SC designed the study. JYD performed statistical analyses and visualization. VP, PG, and SC supervised the study. PG acquired funding. JYD wrote the original draft with contributions from VP, PG, and SC. All authors contributed to data interpretation, manuscript revision, and final approval of the submitted version.

