## Supplementary figures for "Diatoms vs dinoflagellates: a temporal network analysis of bloom impacts on diversity and phytoplankton community structure in French coastal waters"

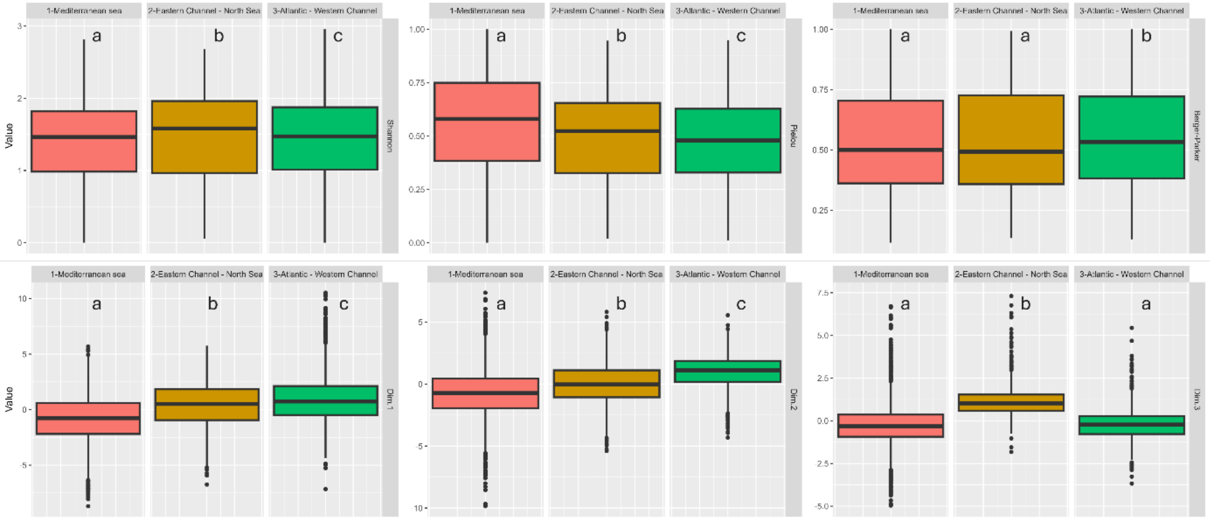


**Supp. Fig. 1: Differences between regions for diversity indices (upper part) and PCA dimensions based on graph metrics (lower part).** Letters indicate significant differences ($p < 0.05$) according to a Kruskal-Wallis test.

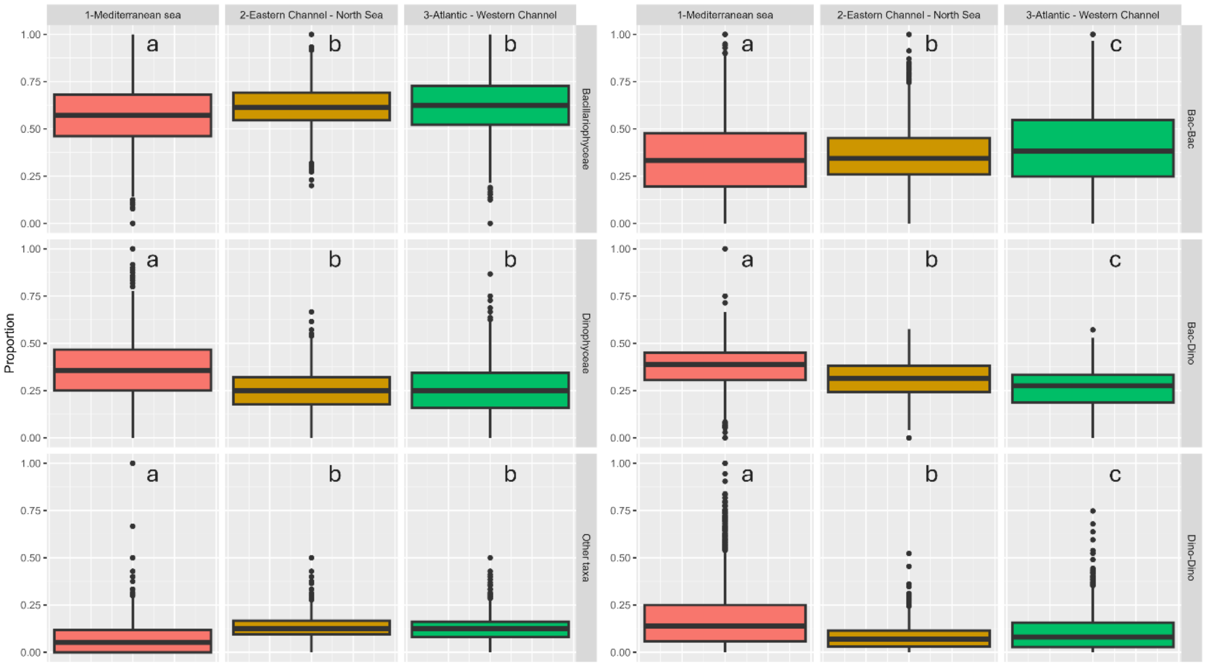


**Supp. Fig. 2: Differences between regions for the taxonomic (A) and association types (B) composition of the temporal association networks.** Letters indicate significant differences ($p < 0.05$) according to a Kruskal-Wallis test.

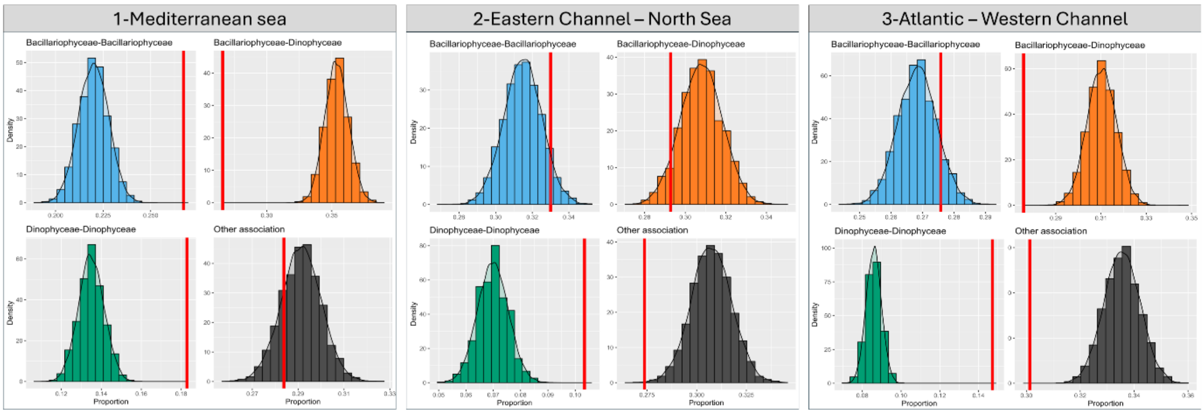


**Supp. Fig. 3: Distribution of each association type proportion obtained with the random association networks (bar plots) and the observed association type proportion (red vertical line) in the global networks for each region. The bar plots are colored according to the association type (between *Bacillariophyceae*, light blue; between *Bacillariophyceae* and *Dinophyceae*, orange; between *Dinophyceae*, dark green; and other types of associations, dark)**

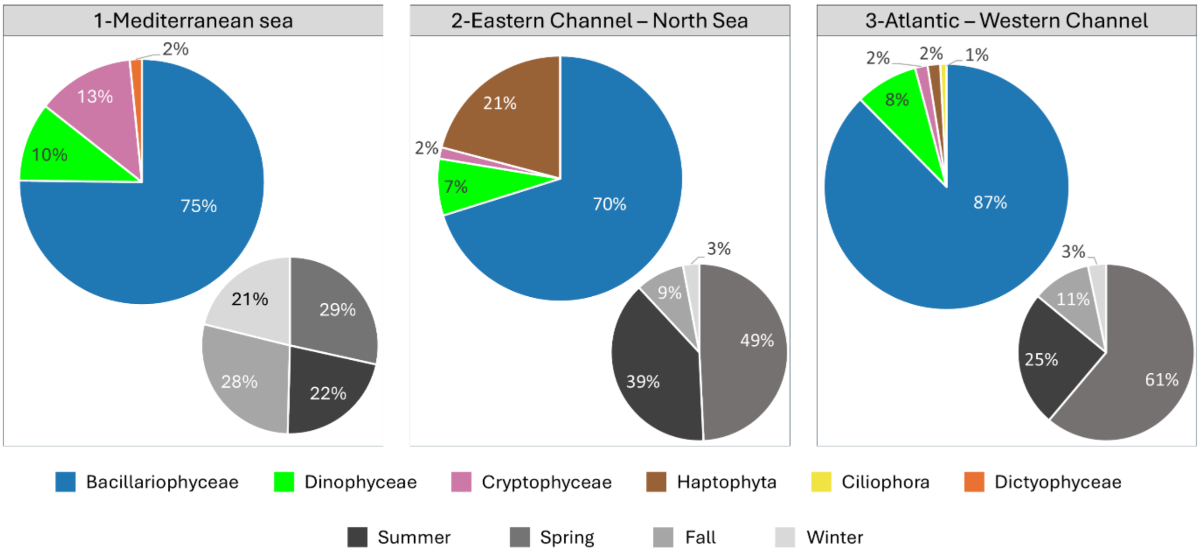


**Supp. Fig. 4: Composition of the detected blooms by region.** The large colored pie chart shows the relative proportion of blooms detected by class, while the smaller grayscale pie chart displays the relative proportion of blooms detected by season.

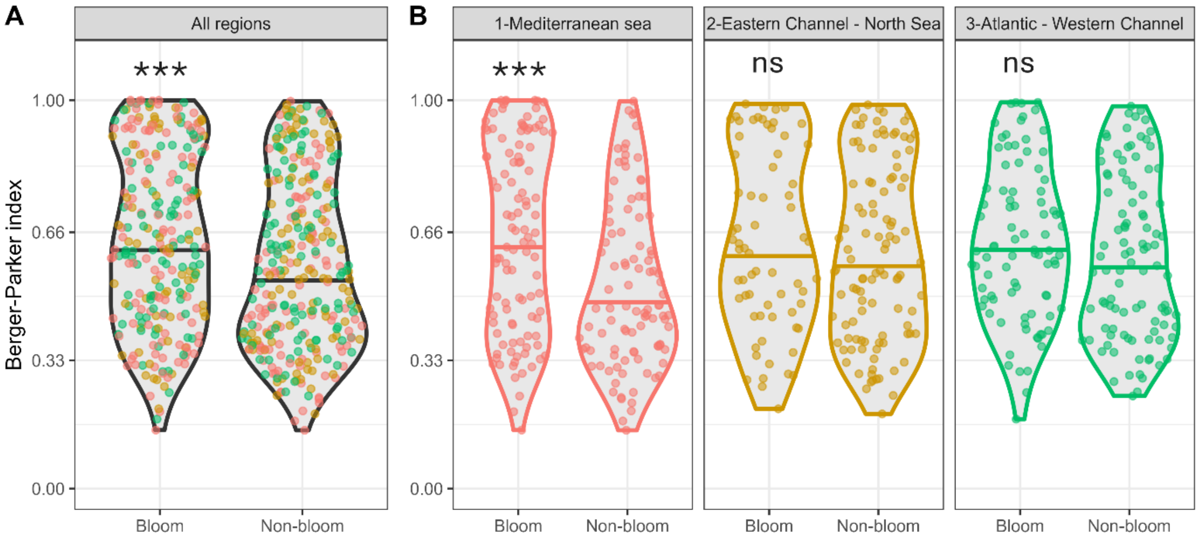


**Supp. Fig. 5: Berger-Parker index during bloom and “non-bloom” sampling for all regions (A) and by region (B).** The horizontal line represents the mean value. Points and violin plots are colored according to the region (Mediterranean Sea, red; Eastern Channel–North Sea, brown; Atlantic–Western Channel, green). Asterisks indicate whether the Berger-Parker index during blooms is significantly different from “non-bloom” periods according to a Wilcoxon test (***: $p < 0.0001$, **: $p < 0.001$, *: $p < 0.05$, ns: $p > 0.05$).
